# Cryo-EM Map of the CFA/I Pilus Rod

**DOI:** 10.1101/586412

**Authors:** Weili Zheng, Magnus Andersson, Narges Mortezaei, Esther Bullitt, Edward H. Egelman

**Affiliations:** Department of Biochemistry & Molecular Genetics, University of Virginia, Charlottesville, VA, USA; Department of Physics, Umeå University, Umeå, Sweden; Department of Physiology & Biophysics, Boston University School of Medicine, Boston, MA, USA

**Keywords:** Fimbriae, bacterial adhesion, helical reconstruction, force spectroscopy, electron cryomicroscopy

## Abstract

Enterotoxigenic *Escherichia coli* (ETEC) are common agents of diarrhea for travelers and a major cause of mortality in children in developing countries. To attach to intestinal cells ETEC express colonization factors, among them CFA/I, which are the most prevalent factors and are the archetypical representative of Class 5 pili. Due to their helical quaternary structure that can be unwound and works as a damper of force, CFA/I pili help ETEC bacteria withstand intestinal fluid motion. We report in this work the CFA/I pilus structure at 4.0 Å resolution and report details of the donor strand complementation. The density map allows us to identify the buried surface area between subunits, and these regions are correlated to quaternary structural stability in class 5 and Chaperone-Usher pili. In addition, from the EM map we also predicted that residue 13 (proline) of the N-terminal β-strand could have a major impact on the filament’s structural stability. Therefore, we used optical tweezers to measure and compare the stability of the quaternary structure of wild type CFA/I and point-mutated CFA/I in which proline 13 was changed to a non-polar residue, phenylalanine. We found that pili with this mutated CFA/I require a lower force to unwind, supporting our hypothesis that Pro 13 is important for structural stability. The high-resolution CFA/I pilus structure presented in this work and the analysis of structural stability will be useful for the development of novel antimicrobial drugs that target the pilus structure to reduce its damping properties, which are needed for initial attachment and sustained adhesion of ETEC.

**Synopsis:** The structure of a common virulence factor expressed on the surface of diarrheacausing bacteria, CFA/I pili, has been determined at 4.0 Å resolution. The role of proline 13 in stabilizing the pilus structure has been confirmed using force-measuring optical tweezers on wild type and point-mutated pili.

## 1. Introduction

Enterotoxigenic Escherichia coli (ETEC) bacteria are common agents of infectious diarrheal diseases affecting hundreds of millions people each year, with hundreds of thousands deaths (Kotloff et al., 2013; Platts-Mills et al., 2015). To infect a host, ETEC first needs to adhere, a step that is mediated by adhesion pili (also called ‘fimbriae’) (Anantha et al., 2004). CFA/I pili, the archetype of class 5 pili, are the most prevalent ETEC colonization factor and extensive evidence indicates that either CFA/I pili or constructs of pilin subunits work as protective antigens (Freedman et al., 1998; Luiz et al., 2015).

CFA/I pili are assembled by the alternate chaperone pathway into helical polymers expressed on the ETEC surface (Soto & Hultgren, 1999). The bioassembly of CFA/I pili is initiated by the attachment of the chaperone/minor pilin (CfaA/CfaE) heterodimer to the usher protein CfaC at the outer membrane assembly site followed by cyclic incorporation of the major pilin, CfaB (Poole et al., 2007; Li et al., 2009). Both pilin subunits, CfaE and CfaB, require the chaperone CfaA for folding and stability and are incorporated into the pilus by the donor-strand exchange mechanism (Sauer, 1999; Choudhury, 1999). At the outer membrane, an usher protein facilitates donor-strand exchange, during which the N-terminus of each pilin subunit is transferred from the chaperone protein to a growing chain of pilin subunits (Choudhury, 1999; Sauer, 1999; Verger et al., 2007). This growing chain of subunits forms a helical structure 1–3 μm in length and with an outer diameter of approximately 8 nm (Mu et al., 2005, 2008).

Similarly to other adhesion pili expressed by ETEC, for example, CS2 and CS20, the quaternary structure of CFA/I can be unwound by tensile force (Andersson et al., 2012; Mortezaei, Singh et al., 2015; Mortezaei, Epler et al., 2015). Tensile force sequentially unwinds subunits in the quaternary structure into a linearized open-coiled structure, thus producing a constant force response. Via physical simulations it has been shown that when a bacterium is exposed to fluid shear forces, the capability to unwind reduces the load on the adhesin expressed at the tip (Zakrisson et al., 2012, 2015). Therefore, bacteria’s ability to attach and stay attached to host surfaces are strongly correlated to the mechanics of pili. Further evidence for this was shown in an in vivo mouse study, in which uropathogenic E. coli bacteria expressing point mutated type 1 pili (also a donor-strand pilus type) that required a reduced force to unwind the quaternary structure were significantly attenuated in bladder infection and intestinal colonization (Spaulding et al., 2018). Thus, understanding the intrinsic interactions and essential properties of pili is important to better understand their role during colonization. However, what intrinsic interactions and essential properties that stabilize the CFA/I pili are not perfectly clear.

We show here the structure of CFA/I pili at 4.0 Å resolution. A fit of the crystal subunit into the map illustrates conformational differences between the known crystal structures of the major pilin subunit, CfaB (Bao et al., 2016) and its structure within an assembled pilus filament. We present intrinsic properties responsible for quaternary structure stability, determined by evaluating the buried surface area of subunit-subunit interactions from the cryo-EM map.

## 2. Results

### 2.1. CFA/I pili structure

Our electron cryomicroscopy (cryo-EM) three-dimensional helical reconstruction of CFA/I pili (Figs 1A, S1) shows pilin subunits assembled into 7.8 nm diameter helical filaments with an 8.6 Å rise per CfaB subunit, and 3.18 CfaB subunits per turn of the helix. After rigid body fitting of the subunit’s structure determined by x-ray crystallography (7) into the cryo-EM helical reconstruction, we used RosettaCM to refine the filament structure of CfaB based upon the cryo-EM map. The fit of the subunit into the cryo-EM map is shown in Figs 1B, C, with a ribbon backbone display and a full amino acid display, respectively. CFA/I pili are assembled using a beta-strand N-terminal extension of the nth subunit fitted into the groove created by a ‘missing beta-strand’ in the previously added n -1st subunit. This interaction provides strong non-covalent bonds between adjacent subunits. As seen in Figs 2A, B, the outer face of the subunit is mostly hydrophilic (Fig. 2A), whereas there is a hydrophobic groove visible when the subunit is rotated 90° (Fig. 2B). This groove gets filled by the β-strand N-terminal extension of the next subunit as the pilus is assembled. Stabilization of the pilus as a helical filament appears to be primarily through contacts between the nth and n+3rd subunits (Fig. 3). Calculated from our CFA/I model using CocoMaps software (Vangone et al., 2011) the buried surface area between subunits n and n+3 is 1,087 Å 2, whereas the buried surface area between n and n+1 is 485 Å 2, and between n and n+2 is 453 Å 2. Subunit-subunit interactions that produce the buried surface area are described in Supplemental Table 1. Once assembled into the helical filament, positively- and negatively-charged surface residues are distributed throughout the pilus outer surface, while many negative charges are exposed on the inner surface, as seen in Figs 2D and 2E, respectively.

**Figure 1:**
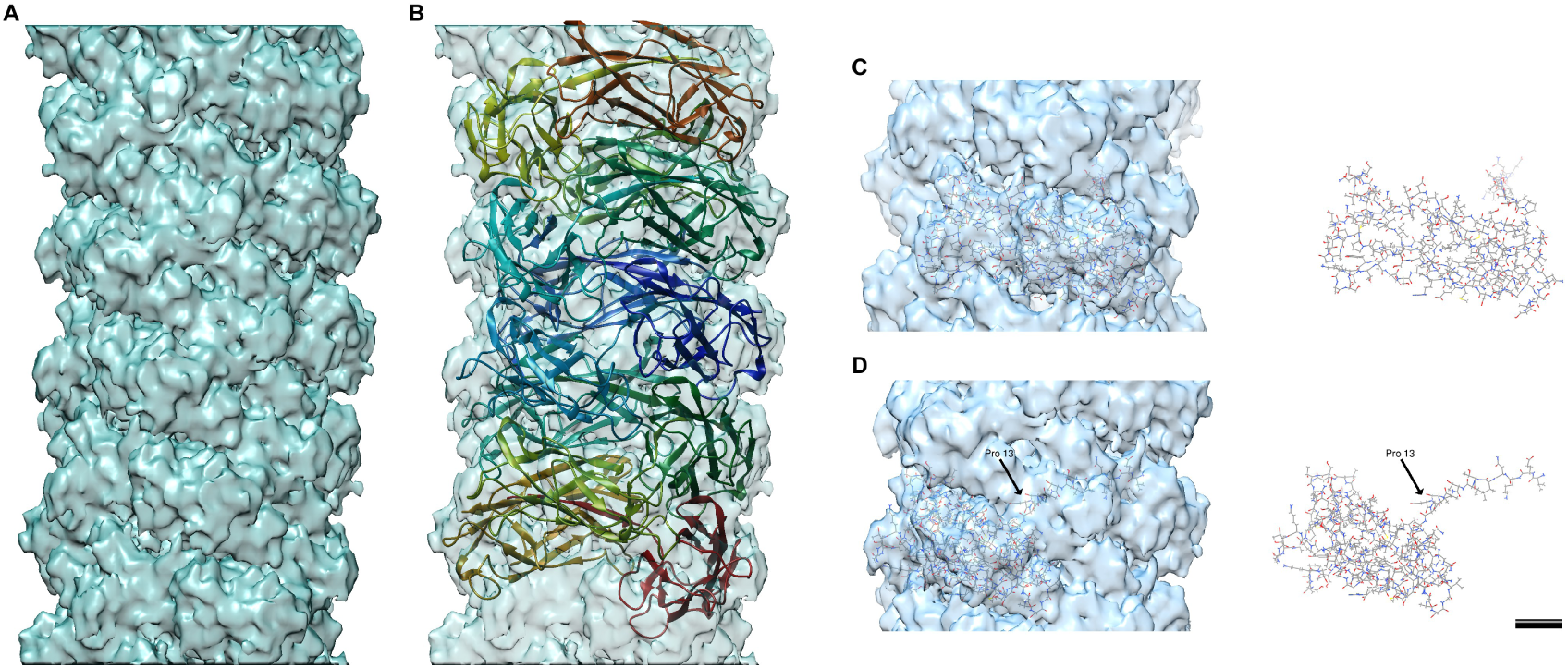
Three dimensional structure of CFA/I pili by cryo-EM. A density map of CFA/I plii (A) has been fitted with CfaB subunits shown as ribbons (B) and atoms (C-D). The pilus in D is rotated 45° about the helical axis from that shown in C, to highlight the position of Pro 13.

**Figure 2:**
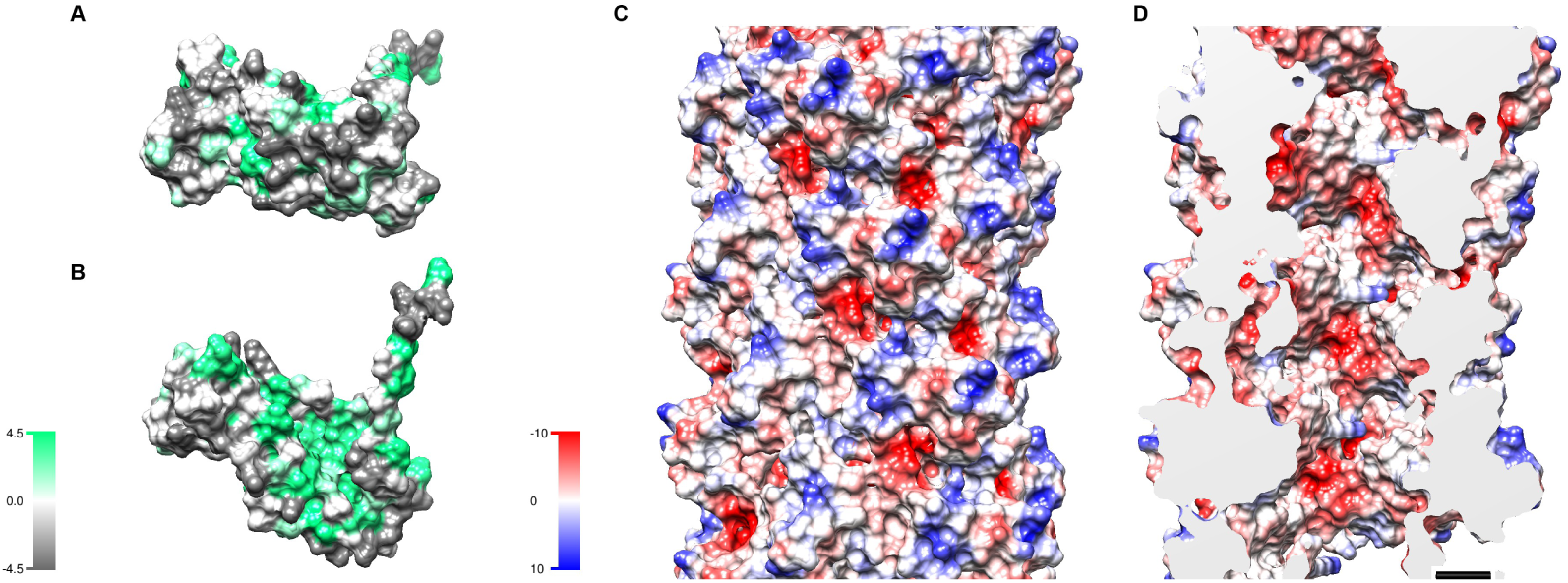
Hydrophobicity and surface charges. Hydrophobicity maps of surface-exposed residues for individual subunits are shown oriented so that the outer surface (A, top) or the surface that accepts the N-terminal extension from the adjacent subunit (B, bottom) is visible. The charge distribution on the outer surface (C) and the inner surface (D) of the pilus shows a negatively charged inner surface that could provide a target for positively charged disruptive therapeutic molecules/peptides such as histatins (Brown et al., 2018).

**Figure 3:**
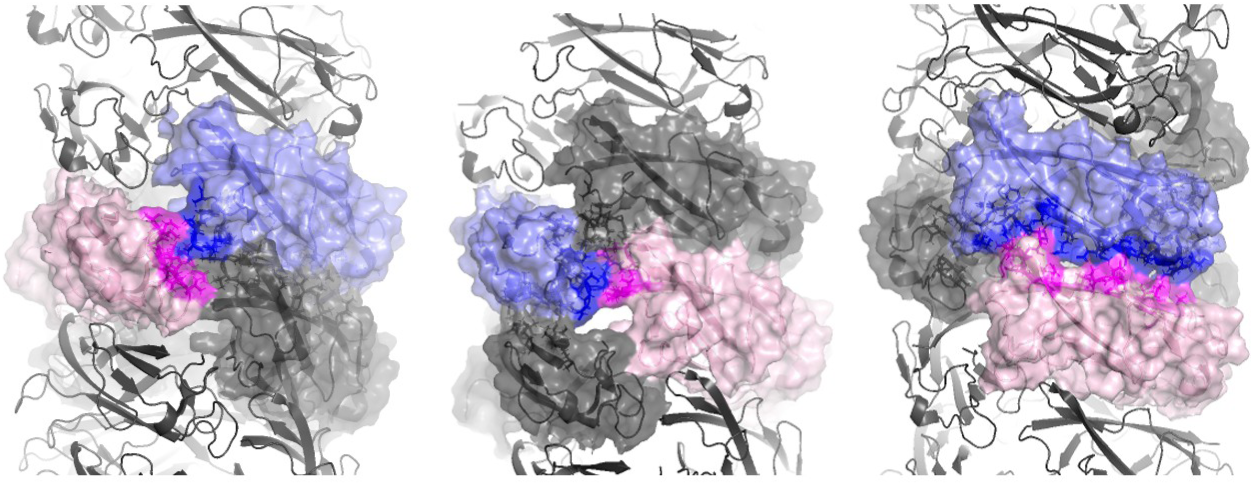
Buried surface area. The majority of the buried surface area between subunits in the assembled CFA/I pilus structure is between subunits n and n+3. Subunit n, blue; contact subunit, pink; buried surface area, magenta.

The most significant conformational change in the structure of CfaB, as compared to a trimer structure (PDB ID: 3F85) determined by x-ray crystallography (7), is an approximately 110° rotation of the N-terminal β-strand. In both structures, 12 N-terminal amino acids, termed the ‘N-terminal extension’ (NTE), have an approximately linear orientation. When expressed as a trimeric oligomer and crystallized, the subsequent amino acids continue in an approximately linear orientation. In contrast, there is an approximately 110° rotation of the NTE observed in our cryo-EM map. This rotation reorients the rest of the subunit, resulting in assembly of CfaB subunits into the helical pilus filament. While the cryo-EM map does not have sufficient resolution to definitively assign orientations of residue side chains, the CfaB backbones are well-aligned after residue 13 (Val 14, Ile 15, Asp 16, etc.). More proximal to the N-terminus, the backbone diverges by 110° at Pro 13, and then continues approximately linearly in this new direction through the N-terminal residue.

Thus, the amino acid Pro 13 appears to be the hinge point for the conformational difference that is observed between the CfaB crystal structure, with its linearly arranged subunits, and the CfaB cryo-EM filament structure (Fig. 4). In the pilus filament structure, there are 13 contacts between Pro 13 of the nth subunit and other CfaB subunits. All observed contacts of Pro 13 are to residues in the previously added n-1st subunit. A visualization of how this movement might occur is shown in the supplemental movie. Additional, smaller, conformational changes between CfaB in its fibrillar state, as compared to the CFA/I helical filament, include a reorientation of a loop from residues 103-109 (Fig. 3 and supplemental movie). In the pilus filament, this loop comes down from the subunit above, occupying part of the location of the ‘staple’ that was seen previously to create an interaction between the n-4th and nth subunits in the P-pilus structure (19). We note that in CFA/I pili there are no contacts between the subunits via this loop. The orientations of two other loops within CfaB also change: residues 33-39, and residues 62-66, with Pro 66 providing a possible hinge region for this loop movement.

**Figure 4:**
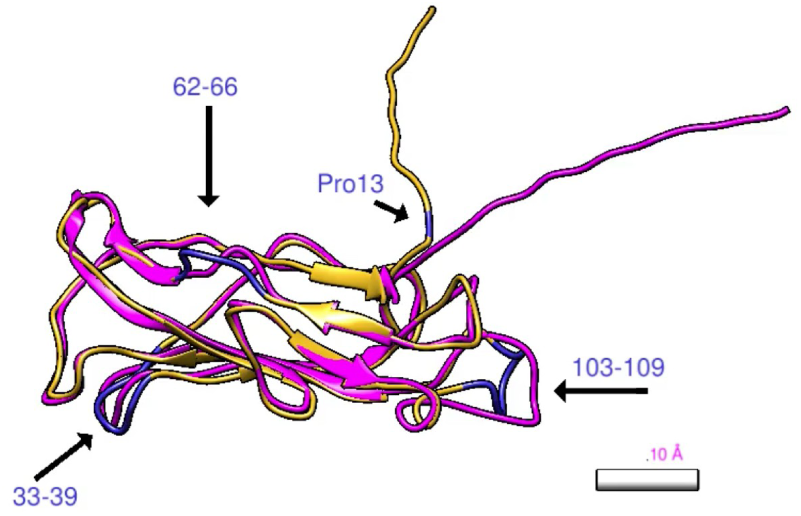
Hinge region difference. The major conformational change between subunits adopting a linear versus helical macromolecular assembly is a rotation of 110° of the N-terminal extension, residues 1-13. Additional smaller differences occur in loops at residues 33-39, 62-66, and 103-109. CfaB central subunit from pdb 3F85, pink; This research, magenta. Scale bar 10 Å.

### 2.2. Pro 13 of CfaB influences quaternary structural stability under tensile force

To investigate the impact of Pro 13 on quaternary structural stability of CFA/I pili, we used optical tweezers to measure the force required to extend individual pili. By pulling on a bead nonspecifically bound to the tip of a CFA/I pilus on a tethered bacterium, we extended the pilus from its helical to unwound shape and thereby assessed its mechanical properties. Thus, we could quantify the force needed to reorientate CfaB subunits.

First, we investigated the mechanical properties of CFA/I pili expressed by wild type bacteria. We show in Fig. 5A a representative force-extension response curve of one such pilus. The data indicate an initial linear increase in force. This initial linear region of constant force indicates stretching of a linearized structure. The quaternary structure then unwinds, represented by the constant force plateau between 0.1 – 3.4 µm, after At 3.8 µm the applied force is so high that the pilus detaches from the bead resulting in the sudden force drop.

**Figure 5:**
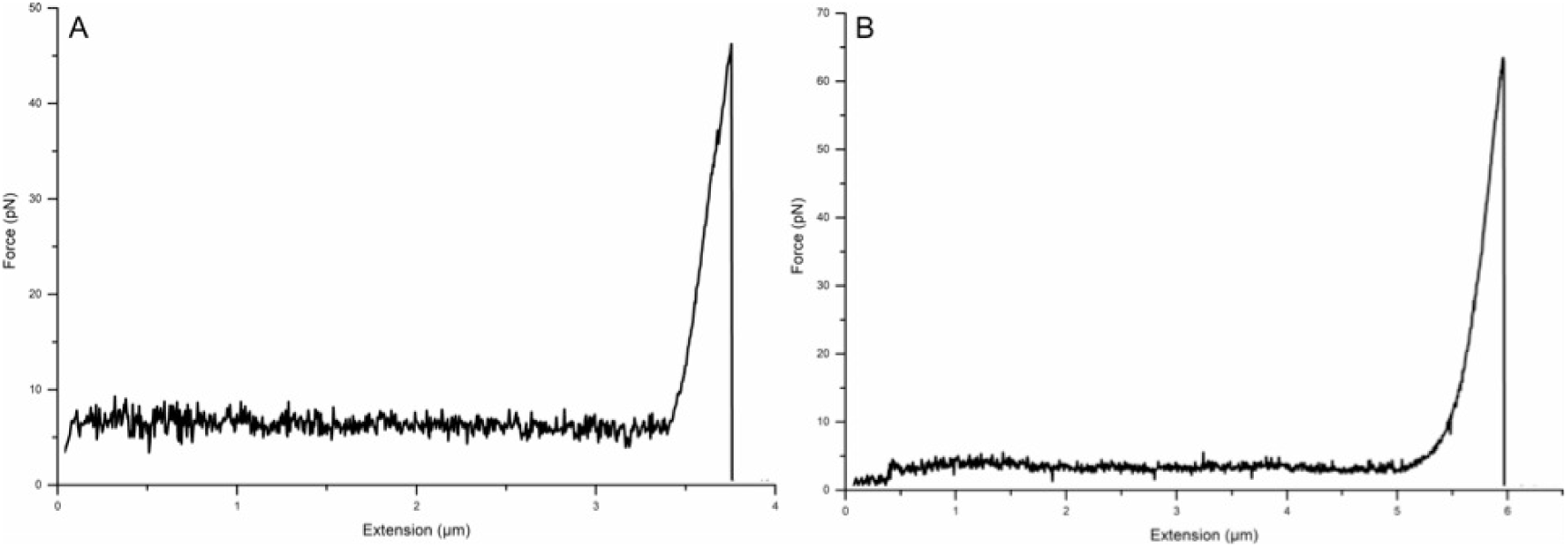
Pro 13 is essential for the quaternary stability of CFA/I pili. Mutation of Pro 13 to phenylalanine in CfaB changed the force required to unwind the quaternary structure of a pilus. (A) Force response of a single wild type CFA/I pilus using optical tweezers. (B) Force response of a single mutated CFA/I pilus.

Since the plateau force is an indicator of quaternary stability, a high unwinding force implies better stability to tensile forces. We averaged the force data of the plateaus for all measured pili, which for the wild type CFA/I was 7.0 ± 1.0 pN (mean ± standard deviation (SD), n = 20, two biological replicates). Thus, these data are in agreement with previous force measurements on CFA/I (Andersson et al., 2012). Next, we investigated the point mutant CFA/I pili in which Pro 13 was changed to a non-polar residue, phenylalanine. Cells expressing these mutated structures have shown a reduced hemagglutination capability when exposed to agitation for short intervals (Li et al., 2009). Also, in that study intact helical filaments were rarely observed. Thus, we hypothesized that force curves should be different than observed for wild type CFA/I.

Indeed, our force measurements showed that this point mutation changed structural stability. Using the same force measuring approach as for the wild type pili we found that the data for mutated pili indicate a structure that is less able to withstand tensile force, Fig. 5B. The average plateau value force was 4.5 ± 1.5 pN (n = 26, five biological replicates). An unpaired, two-sided t-test comparison between the force needed to unwind two types of pili indicates a significant difference (p-value < 0.00001).

## 3. Discussion

As shown previously (Li et al., 2009), the CFA/I pilus filament is assembled via donor strand complementation, where the hydrophobic N-terminal region of an incoming subunit (n+1) fits into an available hydrophobic groove in the nth subunit. In our new higher resolution structure, details of this interaction and the positioning of the β-strands that comprise the subunit are now visualized. The CfaB subunit now fits into the cryo-EM map with a 12° rotation perpendicular to the helical axis, as compared to its fit into a map produced from negatively stained pili (Mu et al., 2008). In addition, the previous structure appeared to have openings to the central cavity, whereas at higher resolution there are grooves that do not extend to the pilus center (Fig. S2).

Our findings suggest that quaternary structural stability for CFA/I pili is due mainly to the stacking interface formed between every n and n+3rd subunit, as demonstrated by the largest buried surface area, 1,087 Å 2. We challenged this interaction by applying tensile force using optical tweezers and measured a 7.0 ± 1.0 pN force needed to unwind CFA/I pili. This finding correlates well with what has been shown for type 1 and P pili as both of these pilus types have more buried area, 1,453 Å 2 and 1,616 Å 2 (Hospenthal et al., 2016, 2017; Spaulding et al., 2018), and require stronger unwinding forces (Andersson et al., 2007). However, there appear to be additional subunit-subunit interactions in CFA/I pili that have a large influence on its quaternary stability. For example, mutations in Pro 13 have been shown to cause significant phenotypic changes in the CFA/I pilus. Analysis of EM images showed that a large majority of pili observed by electron microscopy were 2 nm diameter fibrillar structures, rather than 8 nm diameter filamentous (helical) structures (Li et al., 2009). From our new structural and biophysical data, we expect that this increase in unwound pili is due to the large number of missing contacts that could not be formed between Pro 13 and its contact subunits, n-1 and n+1. Without these contacts, it appears to be difficult to form and maintain a helical filament. Conversely, the linear fibrillar structure would likely be uncompromised, as Pro 13 does not participate in the n to n+1 NTE interaction. These data are validated by our measurement of a lower unwinding force for CFA/I pili in which Pro 13 was mutated to phenylalanine.

The surface charge and hydrophobicity maps of CFA/I pili show characteristic positive, negative, and some hydrophobic residues on the outer surface of the pilus. Of particular interest is the highly negatively charged inner surface of the pilus. Further studies will be used to test a mechanism for disruption of pili by positively charged antimicrobial peptides, such as histatin 5, via interaction with this negatively charged surface (Brown et al., 2018). This detailed view of the structure of CFA/I pili provides an opportunity for developing new strategies to disrupt bacterial adhesion, and limit or prevent subsequent disease.

## 4. Methods

### 4.1. Electron cryomicroscopy

ETEC bacteria E7473 were grown overnight on CFA agar plates (1.0% Casamino Acids, 0.15% yeast extract, 0.005% MgSO4, and 0.0005% MnCl2 2.0% agar) with 50 ug/ml kanamycin at 37°C, overnight. Bacteria were resuspended in 0.5 mM Tris, 75 mM NaCl, pH 7.4 and heated at 65°C for 30 minutes to extract pili. Bacteria without pili were pelleted at 10,000 × g, and the supernatant was centrifuged at 14,500 × g for 30 min to remove cell debris. Pili were precipitated in 300 mM NaCl and 0.1M MgCl2, let sit for 3 ours, and spun at 25,000 × g for 40 min. Pili were resuspended overnight in 0.5 mM Tris, pH 7.4; precipitation and resuspension were repeated and pili were dialyzed against 0.5 mM Tris, pH 7.4.

### 4.2. Cryo-EM data collection and image processing

3 μL of sample was applied to plasma cleaned lacey carbon grids (Ted Pella, Inc., 300 mesh), followed by plunge-freezing using a Vitrobot Mark IV (FEI, Inc). The frozen grids were imaged with a Falcon III direct electron detector (pixel size 1.09 Å /pixel) in a Titan Krios at 300keV. A total of 2,880 movies, each of which was composed of 33 frames with a total dose of ∼45 electrons/Å2, were collected using integration mode. The defocus range was set as 0.5 ∼ 3 μm. Images were motion corrected using MotionCor2 (Zheng et al., 2017), and the program CTFFIND3 (Mindell & Grigorieff, 2003) was used for determining the defocus and astigmatism. Images with good CTF estimation as well as defocus <3μm were selected to use in the following helical reconstruction. The SPIDER software package (Frank et al., 1996) was used for most of the rest operations with the aligned first 15 frames (∼20 electrons/ Å2) of the motion-corrected image stacks. CTF was corrected by multiplying the images with the theoretical CTF, which corrects the phases and improves the signal-to-noise ratio. The e2helixboxer routine within EMAN2 (Tang et al., 2007) was used for boxing the long filaments from the images. A total of 117,011 384 px-long overlapping segments, with a shift of 12 px between adjacent segments (∼97% overlap), were extracted for further use in the IHRSR (Egelman, 2000) reconstruction. With a featureless cylinder as an initial reference, ∼97,295 segments were used in IHRSR cycles until the helical parameters (axial rise of 8.6 A° and rotation of 113.3° per subunit) converged. The resolution of the final reconstruction was determined by the FSC between two independent half maps, generated from two non-overlapping data sets, which was 4 A° at FSC = 0.143 (Fig. S1).

### 4.3. Model building and refinement

We used the previous major pilin CfaB crystal structure (PDB ID: 3F85) as an initial model and docked it into the cryo-EM map by rigid body fitting. Model modification was done by manually editing the model in Chimera (Pettersen et al., 2004) and Coot (Emsley & Cowtan, 2004). We then used the modified model as the starting model for further iterative refinement with RosettaCM (Wang et al., 2015) and Phenix (Adams et al., 2010). The refined protomer model of CfaB was then re-built by RosettaCM with helical symmetry and real-space refined in Phenix to improve the stereochemistry as well as maximize model-map correlation. The final CfaB model was validated with MolProbity (Chen et al., 2010) and the coordinates were deposited in the Protein Data Bank with the accession code 6NRV. The corresponding cryo-EM map was deposited in the EMDB with accession code EMD-0497. The refinement statistics are given in Supplemental Table S2.

### 4.4. Optical tweezers measurements

We carried out force spectroscopy measurements using a high-resolution optical tweezers system constructed around an inverted microscope (Olympus IX71, Olympus) with a high numerical aperture oil-immersion objective (UplanFI 100× N.A. = 1.35; Olympus) (Andersson et al., 2006). Cells and micro-beads are trapped by a single frequency CW diode pumped laser (Cobolt RumbaTM) that provide a Gaussian TEM00 mode with a wavelength of 1064 nm. As force probe, we used surfactant free 2.5 µm white amidine polystyrene latex beads (product no. 3-2600, Invitrogen) which position was monitored by projecting the beam of a low-power fiber-coupled HeNe laser (1137, Uniphase, Manteca) operating at 632.8 nm onto a position sensitive detector (L20-SU9, Sitek Electro Optics, Partille). The detector signal is amplified with preamplifiers (SR640, Stanford Research Systems) and transferred via an analog-digital (A/D) converter card (PCI 6259M, National Instrument) to the computer where quantitative analysis is performed with a customized in-house LabView program. The stability of the setup was prior to the measurements analyzed with Alan-variance method to minimize the noise level (Andersson et al., 2011). We measured the temperature using a thermocouple coupled to the sample chamber to 23.0 °C ± 0.1 °C. In addition, we assumed that the suspension viscosity only varied with temperature, and thus, the viscosity was set to 0.932 ± 0.002 mPa·s.

We suspended bacteria in 1xPBS to a low concentration suitable for single cell analysis. Microspheres were similarly suspended in 1xPBS. We prepared a sample chamber by attaching two pieces of double-sided adhesive tape (product no 34-8509-3289-7, 3M) spaced with 5 mm, on a 24.0 × 60.0 mm coverslip (no.1, Knittel Glass). A 20.0 × 20.0 mm coverslip (no.1, Knittel Glass) was thereafter gently positioned on top of the adhesive tape, thus forming a 5.0 × 20.0 × 0.1 mm flow channel (Mortezaei et al., 2013). We infused the bacterial suspension or the silica microsphere by adding a few µl of suspension at one of the openings allowing capillary forces to fill the chamber. To avoid drying of the sample, we sealed the ends of the chamber by vacuum grease (DOW CORNING®). The sample was thereafter mounted in a sample holder that was fixed to a piezo-stage (Physik Instrument, P-561.3CD stage) in the optical tweezers instrumentation.

## Appendix A. Supplemental Material

### A.1. Supplemental Tables

#### A.1.1. Supplemental Table 1

**Table 1:** Subunit-subunit interactions in the assembled CFA/I pilus filament

| *Interacting subunits* | *# of interacting residues from subunit n* | *# of interacting residues from contact subunit* | *# of hydrophilic-hydrophobic interactions* | *# of hydrophilic-hydrophilic interactions* | *# of hydrophobic-hydrophobic interactions* |
| --- | --- | --- | --- | --- | --- |
| *n to n+1* | 24 | 18 | 53 | 25 | 23 |
| *n to n+2* | 18 | 11 | 31 | 16 | 19 |
| *n to n+3* | 49 | 33 | 99 | 77 | 30 |

#### A.1.2. Supplemental Table 2

**Table 2:** Cryo-EM reconstruction and model refinement statistics

| ***Cryo-EM Data Collection*** |  |
| --- | --- |
| *Microscope* | Titan Krios |
| *Voltage (kV)* | 300 |
| *Camera* | Falcon II |
| *Pixel size (Å)* | 1.09 |
| *Defocus range (μm)* | 0.5~3 |
| ***Helical Reconstruction*** |  |
| *Number of movies* | 2,880 |
| *Number of segments* | 117,011 |
| *Map Resolution (Å)* | 4.0 |
| *Map sharpening B-factor (Å<sup>2</sup>)* | -220 |
| ***Model Refinement and Validation*** |  |
| *MolProbity score* | 1.89 |
| *Clashscore* | 3.52 |
| *Poor rotomers (%)* | 0 |
| ***Ramachandran plot*** |  |
| *Favored (%)* | 88.89 |
| *Allowed (%)* | 11.12 |
| *Outliers (%)* | 0 |
| ***R.m.s. deviations*** |  |
| *Bond lengths (Å)* | 0.006 |
| *Bond angles (°)* | 1.483 |

### A.2. Supplemental Figures

#### A.2.1. Supplemental Figure 1

**Supplemental Figure 1:**
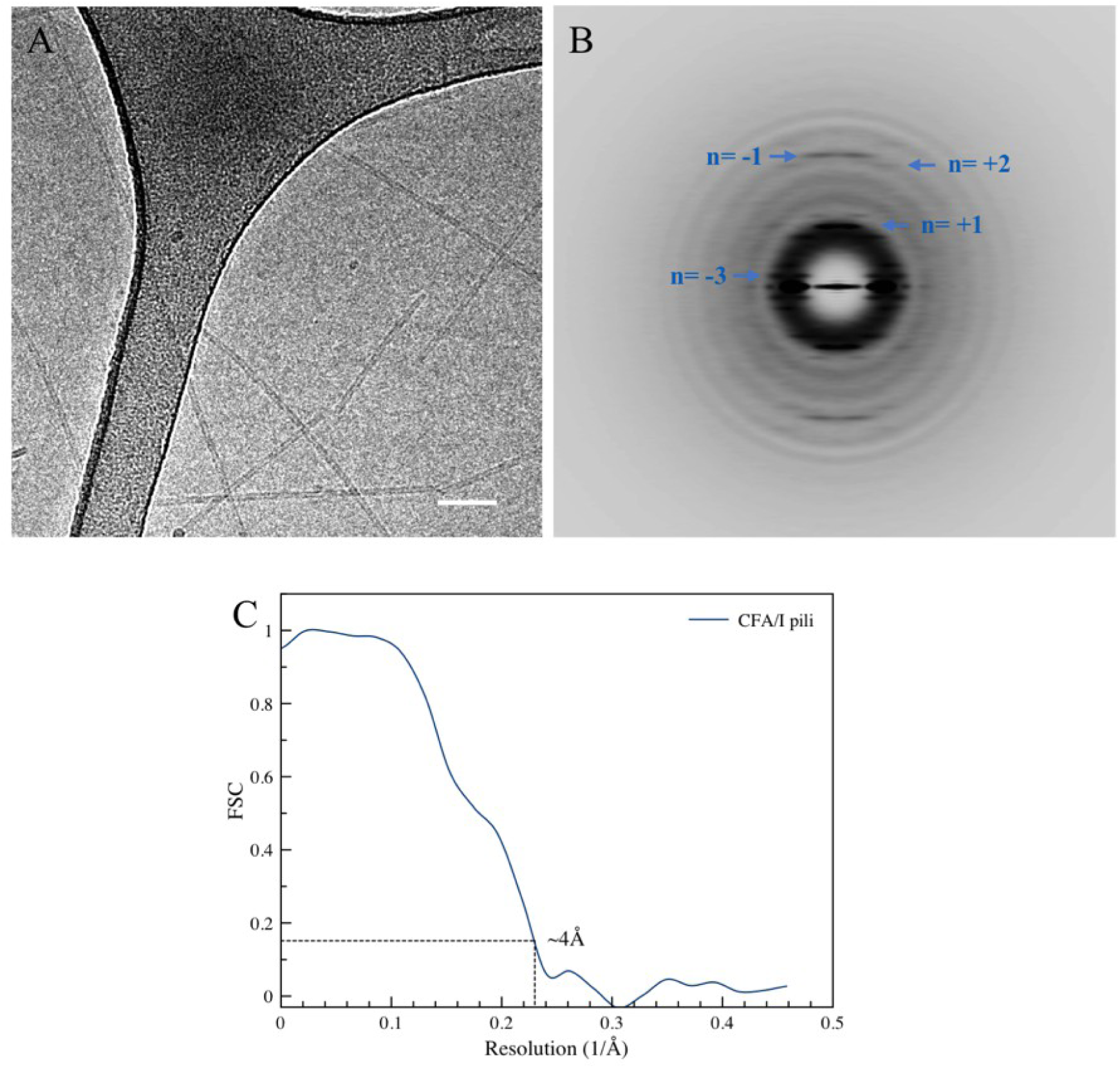
Cryo-EM reconstruction of CFA/I pili. (A) Representative electron micrograph of CFA/I pili in vitreous ice. Scale bar 50nm. (B) The averaged power spectrum, generated from ∼90,000 segments, was used to determine the initial helical symmetry. The Bessel orders are labeled for a number of layer lines. (C) The FSC between two-independent half maps, each generated from non-overlapping datasets, shows a resolution of 4 Å at FSC=0.143.

#### A.2.2. Supplemental Figure 2

**Supplemental Figure 2:**
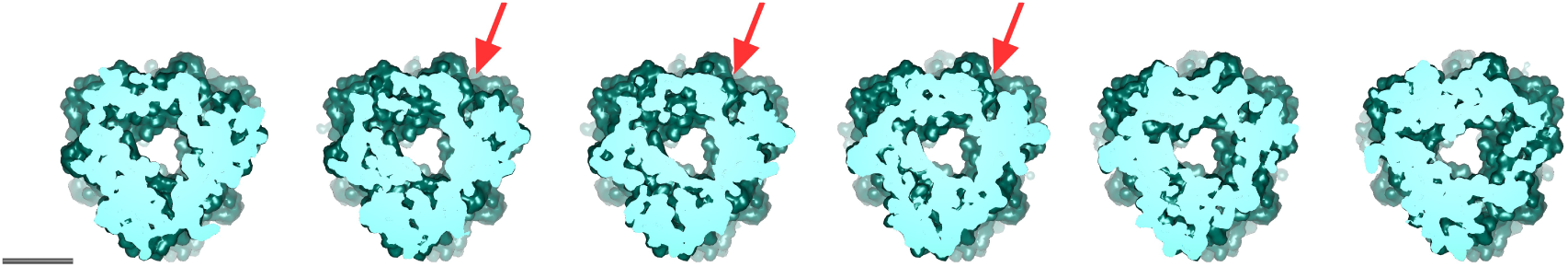
Cross-sectional views of CFA/I pili. Sections every 2 Å showing a groove in the pilus structure without an opening to the central cavity. Scale bar 25 Å.

## Appendix B. Supplemental Movie

Movie shows a visualization of how rotation about Pro 13 could re-orient the direction of a subunit to coil into the helical CFA/I pilus filament.

